# MIF Regulates M1 Macrophage Polarization via CD74/CXCR2/JNK Pathway and Mediates Aortic Dissection in Mice

**DOI:** 10.1101/2023.10.26.564292

**Authors:** Lu Wang, Huishan Wang, Liming Yu, Hui Jiang, Lin Xia

**Affiliations:** General Hospital of Northern Theater Command; Shenyang Northern Hospital

**Keywords:** MIF, aortic dissection, macrophage polarization, JNK, co-culture, VSMC phenotype switch

## Abstract

**Background:** Macrophage polarization and vascular smooth muscle cell (VSMC) phenotypic switching are important features and critical targets in the progression of Aortic dissection (AD). High expression of macrophage migration inhibitory factor (MIF) in aortic and blood specimens has been observed in patients with aortic dissection, but its precise function and mechanism in AD are unknown. We aimed to clarify whether MIF mediates the development of aortic dissection via modulation of M1 macrophage polarization and its specific regulatory pathways.

**Methods:** Based on the BAPN/Ang II-induced acute aortic dissection model and by intraperitoneal injection of the MIF antagonist ISO-1 to inhibit MIF activity in mice. We assayed macrophage infiltration, polarization, and VSMC phenotypic switching in the aorta of mice in each group. Further, we evaluated the polarizing effects of MIF on RAW264.7 cells directly or indirectly in vitro experiments and explored the specific cellular signaling pathways that mediate its function. At last, we evaluated the role and possible mechanisms of RAW264.7 cells with different degrees of polarization in inducing phenotypic switching of MOVAS cells based on a model of indirect co-culture of RAW264.7 cells with MOVAS cells.

**Results:** Pharmacological inhibition of MIF decreased the incidence of BAPN/Ang II-induced aortic dissection and attenuated aortic vascular remodeling in mice by reducing M1 macrophage infiltration in mouse aorta. Through in vitro assays, we demonstrated that MIF could activate the intracellular JNK/c-Jun signaling pathway by targeting the CD74/CXCR2 receptor, promote M1 polarization and upregulate the expression of the M1 macrophage markers, iNOS, IL-18, and CD86 in RAW264.7 cells. Further experiments confirmed that upon co-culture with MIF-induced M1 macrophages, the NF-κB pathway was activated in MOVAS cells, inducing the onset of phenotypic switching and apoptosis.

**Conclusions:** The results indicated that MIF mediated macrophage polarization and regulated the progression of aortic dissection, which provided new scientific evidence for the pathogenesis of aortic dissection, and also suggested that MIF may be a potential preventive and therapeutic target for aortic dissection and aortic-related diseases.

## Introduction

Aortic dissection (AD) is an extremely dangerous and complex cardiovascular emergency with an insidious onset and rapid progression, which can lead to serious complications at early stage due to malperfusion of vital organs and even death due to aortic rupture.^1^ The annual incidence of AD is about 3/100,000,^2,3^ and the prevalence of AD has been increasing in recent years.^4^ Although emergency surgical interventions have significantly improved survival rates in acute type A aortic dissection, surgical mortality remains high.^5^ Therefore, current research focuses on the pathogenesis of aortic dissection as a way to inhibit or control the occurrence and progression of aortic dissection as a potential therapeutic strategy.

Inflammatory cells are closely associated with vascular smooth muscle cell (VSMC) apoptosis, phenotypic switch, and extracellular matrix degradation.^6^ Studies on human aortic specimens and mouse models have shown massive aggregation of macrophages at the site of aortic tears.^7,8^ Peripheral macrophages accumulate towards the aortic wall and M0 macrophages polarize into pro-inflammatory M1 macrophages under the effect of cytokines such as Ang II, GM-CSF and LPS.^9,10^ M1 macrophages further degrade the extracellular matrix by secreting MMPs, pro-inflammatory factors such as IL-2, IL-1β, MCP-1, IFN-γ, leading to the AD occurrence.^11^ In contrast, M2 macrophages are involved in the process of aortic repair by inhibiting the production of inflammatory factors and promoting extracellular matrix neogenesis.^12^ Due to its high plasticity, macrophages have the ability to respond to new environment and polarize again even after polarization to a phenotype.^13^

VSMCs, together with matrix components such as elastic fibers and collagen fibers, constitute the mid-layer of the aortic wall.^14^ The differentiated mature contractile VSMCs are responsible for maintaining normal aortic wall integrity, compliance and stress resistance, and still retain their phenotypic regulatory capacity.^15^ The process of dedifferentiation of contractile VSMC to synthetic VSMC under different pathological stimuli is called VSMC phenotypic switch.^16^ Synthetic VSMC have significantly increased proliferation and migration capacity, and they can express new phenotype-specific genes, such as OPN, and secrete extracellular matrix (ECM) components and MMPs that contribute to vascular remodeling.^17^ Therefore, the investigation of mechanisms of macrophage polarization and VSMC phenotypic switch, as well as their causal role in the development of AD, is crucial for understanding the complex pathogenesis of AD.

Macrophage migration inhibitory factor (MIF) is a structurally unique pleiotropic cytokine with enzymatic, chemokine and hormonal properties that is involved in innate and acquired immune responses.^18^ As an important pro-inflammatory factor, MIF has been shown to be associated with the pathogenesis of various inflammatory diseases and tumors by regulating macrophage polarization.^19–21^ There are few researches related to MIF and AD. Differential protein studies of thoracic aortic dissection and normal aortic tissue showed high expression of MIF.^22^ Similar results were seen in abdominal aortic aneurysm specimens, the expression of MIF and MMPs was enhanced in ruptured abdominal aortic aneurysm specimens compared to stable abdominal aortic aneurysm and normal aortic specimens.^23^ These results suggest the possibility of MIF as a biomarker for the identification of AD and assessment of prognosis. Therefore, we believe that a connection exists between MIF and aortic coarctation, but there is no relevant in-depth study.

Based on the pathogenesis of AD and the biological function of MIF, we hypothesized that MIF may affect the development of AD by regulating macrophage polarization. In this study, we investigated the effect of MIF in AD based on the BAPN/Ang II-induced acute aortic dissection model in mice and further clarified the role of MIF in mediating macrophage polarization. We finally identified the effect of M1 macrophages on VSMC phenotypic switch through co-culture model. These findings provide new therapeutic targets and strategies for the prevention and treatment of AD.

## Materials and Methods

### Animals

Sixty 3-week-old male C57BL/6J mice (12-14g), purchased from Hua Fukang Biological Technology Co., Ltd (China), housed under SPF conditions. All mice experiments were approved by the Animal Experimentation Ethics Committee and conducted according to the experimental guidelines.

The acute aortic dissection model in mice was established by continuous feeding with distilled water containing 0.5% BAPN (Sigma-Aldrich, USA) for 2 weeks and subcutaneous injection of Angiotensin II (Abmole, USA) twice on day 15. We used the MIF antagonist ISO-1 (Abmole, USA) to inhibit MIF expression level in mice.

60 mice were randomly divided into three groups: (a) control group (n=20), fed with normal diet, daily intraperitoneal injection of 4% DMSO and subcutaneous injection of saline on day 15; (b) BAPN/Ang II group (n=20), fed with water containing 0.5% BAPN, daily intraperitoneal injection of 4% DMSO, and subcutaneous injection of Ang II (0.72mg/kg) twice on day 15; (c) ISO-1 group (n=20), fed with water containing 0.5% BAPN, intraperitoneal injection of ISO-1 (10 mg/kg/d), and subcutaneous injection of Ang II twice on day 15.

During the modeling period, mice were autopsied immediately after death to determine whether the cause of death was aortic dissection based on the presence of blood clots in the thoracic, pericardial or abdominal cavities. The mice were euthanized on day 16 with an overdose of anesthetic. After autopsy, the overall morphology of the aorta was observed and recorded under stereomicroscope, and blood specimens and all aortic tissue specimens from heart to the iliac artery were collected. The diameter of each segment of the aorta was measured, and the rate of dissection and rupture was calculated.

### Aortic ultrasonography

Ultrasonography of the thoracic aorta was performed on day 7 and day 14 of modeling. Mice were anesthetized by isoflurane inhalation and fixed in a supine position on the operating table, which was tilted 15-30 degrees to the right and adjusted to the aortic arch view. The maximum internal diameter of the ascending aorta, aortic arch and thoracic descending aorta was recorded.

### Histological analysis

Aortic tissues were fixed in 4% paraformaldehyde for 24 hours, then paraffin-embedded after dehydration and transparency steps, and sliced into 4-μm sections. The sections were stained with hematoxylin-eosin (H&E) and elastica van gieseon (EVG) staining. Elastin degradation was scored as follows: grade 1, intact elastin middle layer; grade 2, mild elastin fracture; grade 3, severe elastin fracture; grade 4, complete elastin fracture with visible rupture.

### Immunohistochemical staining

Aortic sections were sequentially processed for antigen repair, inhibition of endogenous peroxidase, and blocking with 10% goat serum. Overnight incubation was performed at 4 °C with the following antibodies, anti-MIF (1:200, ab65869, Abcam), anti-CD86 (1:500, 28058-1-AP, Proteintech), anti-iNOS (1:500, 18985-1-AP, Proteintech) and anti-α-SMA (1:500, #19245, CST). Immunohistochemistry kits were used for secondary antibody incubation and DAB staining, and hematoxylin was used to counterstain nuclear. Ten fields of view were randomly selected under a 40x objective lens and the mean absorbance was analyzed by Image Pro Plus.

### Western Blot analysis

The whole aortic or cellular protein was extracted using RIPA lysis buffer containing protease inhibitors and phosphatase inhibitors. Protein concentration was measured using the BCA kit and protein denaturation was completed. Equal amounts of total protein were separated using 6%-15% SDS-PAGE gel electrophoresis and transferred to PVDF membranes. Membranes were blocked with TBST containing 5% skim milk or 5% BSA for 2 hours and incubated overnight at 4°C with the following antibodies: anti-MIF (1:200, ab65869, Abcam), anti-iNOS (1:500, 18985-1-AP, Proteintech), anti-IL-18 (1:200, #57058, CST), anti Arg-1 (1:1000, #93668, CST), anti-SM22α (1:1000, ab10135, Abcam), anti-OPN (1:100, sc-21742, Santa Cruz), anti-p-JNK (1:200, sc-6254, Santa Cruz), anti-JNK (1:200, sc-7345, Santa Cruz), anti-p-c-Jun (1:100, sc-822, Santa Cruz), anti-c-Jun (1:1000, #9165, CST), anti-CD74 (1:200, sc-6262, Santa Cruz). After washing in TBST, the corresponding secondary antibodies were incubated for 2 hours at room temperature, imaged under a chemiluminescent detection system, and analyzed semi-quantitatively in grayscale using Image J.

### RT-qPCR analysis

Total RNA was extracted from aortic tissue and cells using RNA extraction. RNA was reverse transcribed into cDNA using SweScript RT First Strand cDNA Synthesis Kit, and the obtained cDNA was amplified by PCR using SYBR Green qPCR Master Mix. GAPDH mRNA was used as an internal reference and the relative expression of each gene was calculated using the ΔΔCt method.

### Gelatin Zymographic analysis

According to the gelatin zymography analysis kit instruction, protein samples were separated by non-denaturing electrophoresis in 10% SDS-PAGE gel containing gelatin. After the gels were denaturing and incubated, they were stained using Coomassie Brilliant Blue, and the part of the gel that was degraded by MMPs was left unstained and appeared as a white, nearly transparent area. MMP-2 and MMP-9 positions were determined according to the protein Marker, and grayscale quantitative analysis was performed using Image J.

### ELISA

According to the experimental protocol provided in the mouse ELISA kit, the concentrations of cytokines (IL-6, IL-1β, IL10 and TNF-α) in mouse serum and cell culture media were determined by ELISA.

### Cell Culture and Treatment

Mouse macrophage line RAW264.7 cells was purchased from Oricell^TM^ (Cyagen Biosciences, China), and primary mouse aortic vascular smooth muscle (MOVAS) cells were obtained from the aorta of C57BL/6J mice. Cells were cultured in high-glucose DMEM with 10% FBS, 1% penicillin-streptomycin and incubated in a CO2 incubator at 5% CO2 and 37°C. All cells used in this experiment were between passages 3 to 8.

100ng/ml LPS (Sigma-Aldrich, USA) and 20ng/ml IFN-γ (Novoprotein, China) were used to stimulate RAW264.7 cells as a positive control group for M1 macrophages. Macrophages were stimulated with rMIF (MCE, USA) at concentrations of 10, 50, 100 and 500 ng/ml for 15 min, 30 min, 1h, 6h, 12h and 24h, and cells were collected for relevant experiments. The following reagents were used to give cell pretreatment, MIF inhibitor ISO-1, CXCR2 inhibitor SB225002 (Abmole, USA), CXCR4 inhibitor AMD3100 (Abmole, USA), and JNK inhibitor SP600125 (Abmole, USA).

### Cell Transfection

RNA interference was applied to down-regulate CD74 expression in macrophages. siRNA was designed and synthesized by BIONEER (Korea). Cells were seeded into 6-well plates 24 h prior to transfection, 50 nM siRNA and 8 μL INTERFERin (Polyplus, France) were mixed in Opti-MEM (Gibco, USA), incubated for 10 min at room temperature and added to the cells. WB assayed transfection efficiency or gave further treatment after 24 h of transfection.

### Immunofluorescence staining

Cells were inoculated onto cell climbing slices, cultured in 24-well plates, given the corresponding treatment and then fixed in 4% paraformaldehyde for 15 min. The slices were then permeabilized in PBS containing 0.2% Trition X-100 for 20 min, followed by blocking with PBS containing 5% BSA for 1 h. Cells were incubated with anti-iNOS (1:100) overnight at 4°C in dark, followed by incubation with Alexa Fluor 488 (1:200) for 1 hour at room temperature. Finally, slices were incubated with DAPI for 30 min at room temperature, observed and recorded using confocal microscopy, and fluorescence intensity was analyzed using Image J.

### Flow Cytometric analysis of Macrophage Polarization

Each group of RAW264.7 cells was treated and collected in 1*10^7^/ml cell suspension. The cell samples were incubated with anti-F4/80-FITC (11-4801-82, eBioscience), anti-CD86-PE (12-0862-82, eBioscience) and anti-CD206-APC (17-2061-82, eBioscience) at 4°C for 1 h in dark washed twice with 1 ml FACS buffer and resuspended cells in 500 μl FACS buffer. Samples were assayed using BD FACSCelesta flow cytometer and analyzed by FlowJo. The CD86^+^/CD206^-^ cells were identified as M1 macrophages.

### Co-culture system

The non-contacting co-culture system of RAW26zX-4.7 cells and MOVAS cells was established using the cell chamber insert (0.4 μm pore-size) in six-well plate. First, 1*10^6^ RAW264.7 cells were inoculated in 6-well plates and stimulated by MIF or LPS/IFN-γ for 24h, while 2.5*10^5^ MOVAS cells were inoculated into the cell chambers, incubated overnight and then starved for 24 hours. Cell chambers were inoculated into 6-well plates with RAW264.7 cells, replaced with fresh complete medium, and co-cultured for 48 hours to obtain MOVAS cells within the chambers for further experiments.

### Flow Cytometric analysis of Apoptosis

Apoptosis detection was performed using the Annexin V-FITC Apoptosis Detection Kit. MOVAS cell suspension containing 1*10^5^ cells after co-culture was prepared and resuspended in 195 μl Annexin V-FITC binding buffer. 5 μl Annexin V-FITC and 10 μl PI were added and incubated at room temperature in dark for 20 min. Samples were assayed using BD FACSCelesta flow cytometer and analyzed by FlowJo. Annexin V^+^/PI^-^ cells were identified as early apoptotic cells.

## Results

### 1. ISO-1 attenuates BAPN/Ang II-induced aortic dissection formation in mice

To clarify the role of MIF in the development of aortic dissection, we used BAPN combined with angiotensin II to establish a mouse AD model and gave continuous intraperitoneal injection of ISO-1 for pharmacological inhibition of MIF. The results found that the combination of BAPN/Ang II significantly increased the incidence of AD and the rupture rate (Fig.1C), whereas ISO-1 treatment reduced the prevalence from 90% (18/20) to 55% (11/20) and the rupture rate from 45% (9/20) to 25% (5/20), and no AD formation was observed in the control group (Fig.1A). Both the outer diameter of the isolated specimen and the inner diameter of the aorta suggest that BAPN/Ang II administration caused progressive dilatation in all segments of the aorta, whereas ISO-1 slowed the progression of dilatation, especially in the ascending and descending aorta (Fig.1D-E).

**Figure 1.**
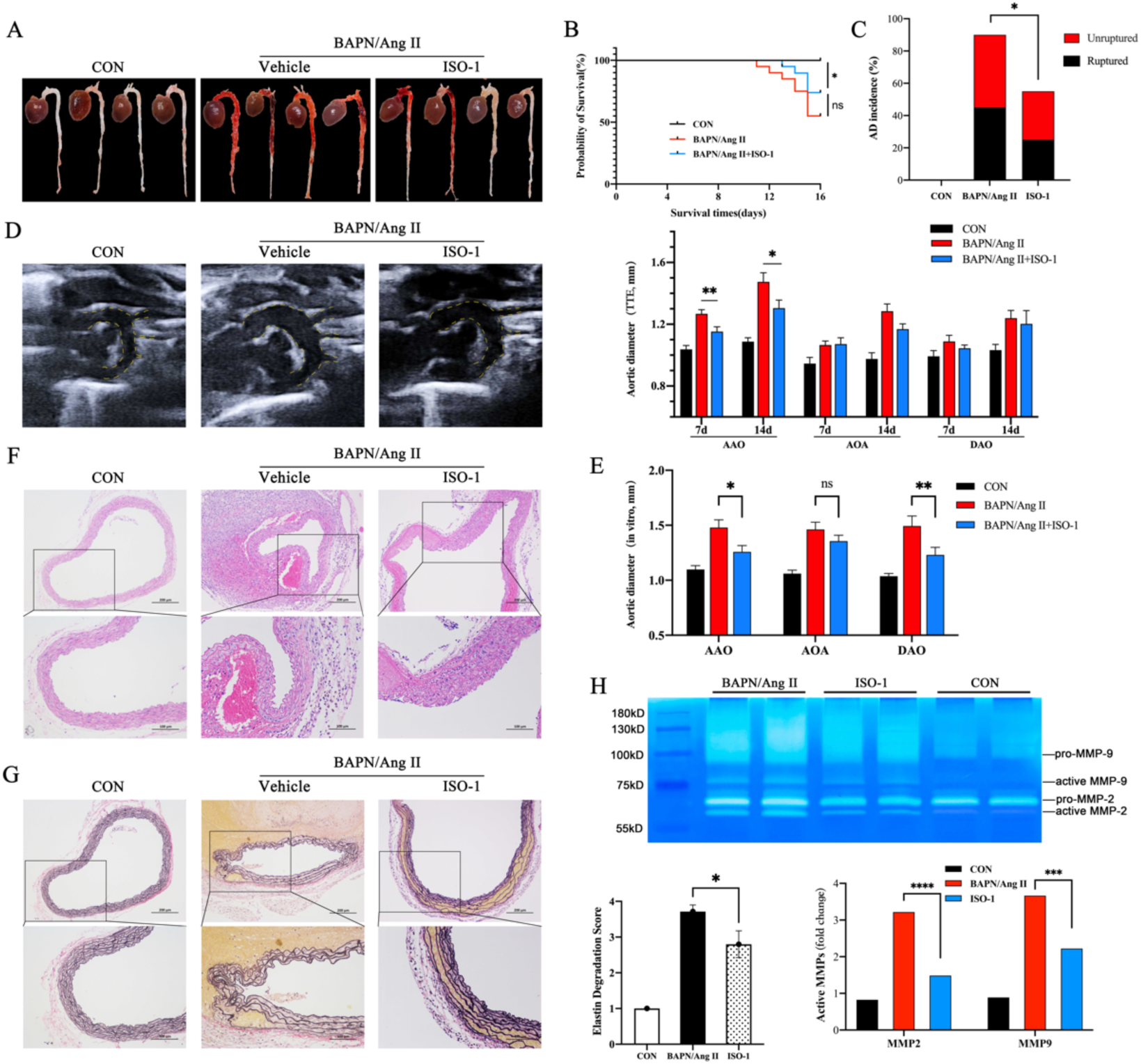
ISO-1 reduces the incidence of BAPN/Ang II-induced aortic dissection in mice. **A**, Representative macrophages of mouse aortic samples from each group. **B**, Kaplan-Meier survival curves in indicated groups. **C**, Incidence and rupture rate of AD. **D**, Representative ultrasound images of the thoracic aorta on day 14 and maximum diameter measurements of each segment. **E**, Maximum diameter of each segment of isolated aortic specimens. **F**, Representative images of HE staining of aortic sections. **G**, Representative graphs of EVG staining of aortic sections, and aortic elastic fiber degradation scores from each group. **H**, Gelatin gel zymography of MMPs activity in aortic tissue.

We clarified the remodeling of the aortic wall in each group by H&E staining and EVG staining (Fig.1F-G). Typical features of aortic dissection such as luminal dilatation, outer membrane thickening, pseudolumen formation, inflammatory infiltration and mid-layer degradation were observed in the sections of AD. In the ISO-1 group, elastic fiber fracture and degradation in the middle aortic layer were alleviated compared to the BAPN/Ang II group, and the elastic fiber degradation score was lower than that of the BAPN/Ang II group. We also assessed the activity of MMPs in aortic tissues by gelatin zymography (Fig.1H), and the results suggested that MMPs activity was significantly higher in the two AD groups compared with the control group, whereas it was lower in the ISO-1 group than in the BAPN/Ang II group (P<0.05).

### 2. ISO-1 inhibits M1 macrophage infiltration in the aorta

Firstly, we assessed the expression of MIF by IHC (Fig.2A), and it was observed that MIF was significantly highly expressed in the peripheral adipose tissue of the diseased aortic wall and in the severely degraded middle layer, and the expression was remarkably higher in the BAPN/Ang II group than in the ISO-1 group (P<0.01).

**Figure 2.**
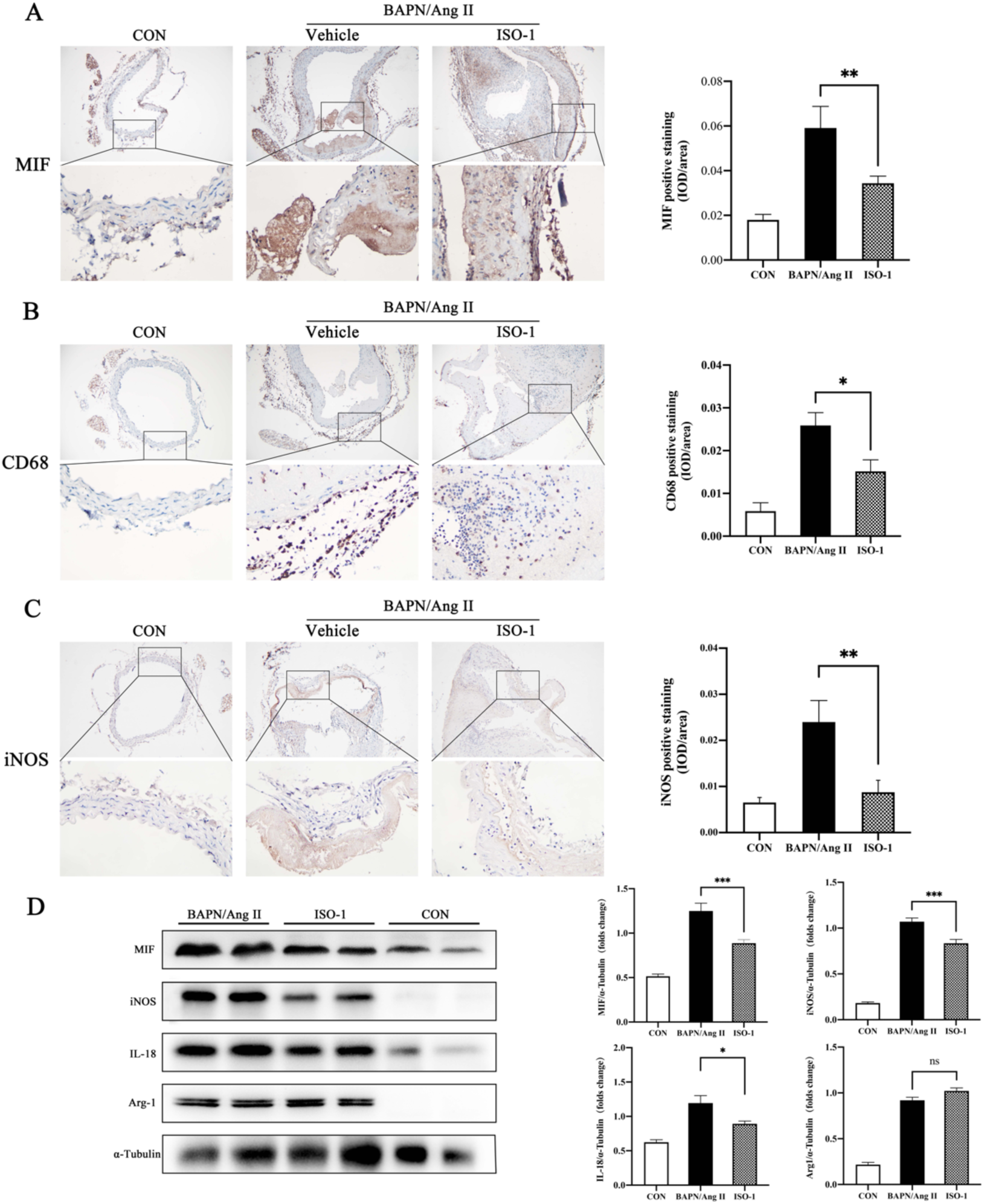
ISO-1 reduces BAPN/Ang II-induced M1 macrophage infiltration in the aortic wall. **A**-**C**, Representative images and quantification of immunohistochemical staining for MIF (A), CD68 (B) and iNOS (C) in aortic sections. **D**, Western blot images and quantification of MIF, iNOS, IL-18 and Arg-1 in aortic tissues of each group. n=5-7; *P<0.05, **P<0.01, ***P<0.001.

To further investigate the infiltration and polarization of macrophages in the aorta, we performed IHC assays for the macrophage marker CD68 and M1 macrophage marker iNOS, respectively (Fig.2B-C). The expression of both markers was significantly elevated in the AD group. Among them, CD68 and iNOS were highly expressed in the outer aortic membrane, perivascular adipose tissue and severely dissected middle layer, which was highly similar to the location of MIF expression. Comparison between groups revealed that the expression of CD86 and iNOS was lower in the ISO-1 group than in the BAPN/Ang II group (P<0.01).

Similar results were verified in Western blot and Rt-qPCR experiments, where the protein expression of MIF and M1 macrophage markers (iNOS, IL-18) were significantly lower in the ISO-1 group than in the BAPN/Ang II group, and the M2 macrophage marker Arg-1 was not significantly different between these two groups (Fig.2D). The mRNA levels of iNOS, IL-6 and TNF-α were significantly increased in the AD group, and the expression of all these mRNAs was decreased in the ISO-1 group compared with the BAPN/Ang II group (Fig.3A). By detecting the relevant cytokines in the serum of each group of mice by ELISA, we found that the concentration of pro-inflammatory cytokine IL-6 was highest in the BAPN/Ang II group and was significantly higher than that in the ISO-1 group (P<0.01) (Fig.3B). Through the above assays, we found that ISO-1 attenuated M1 macrophage infiltration and polarization within the aortic wall and reduced the secretion of pro-inflammatory factors such as IL-6 and TNF-α.

**Figure 3.**
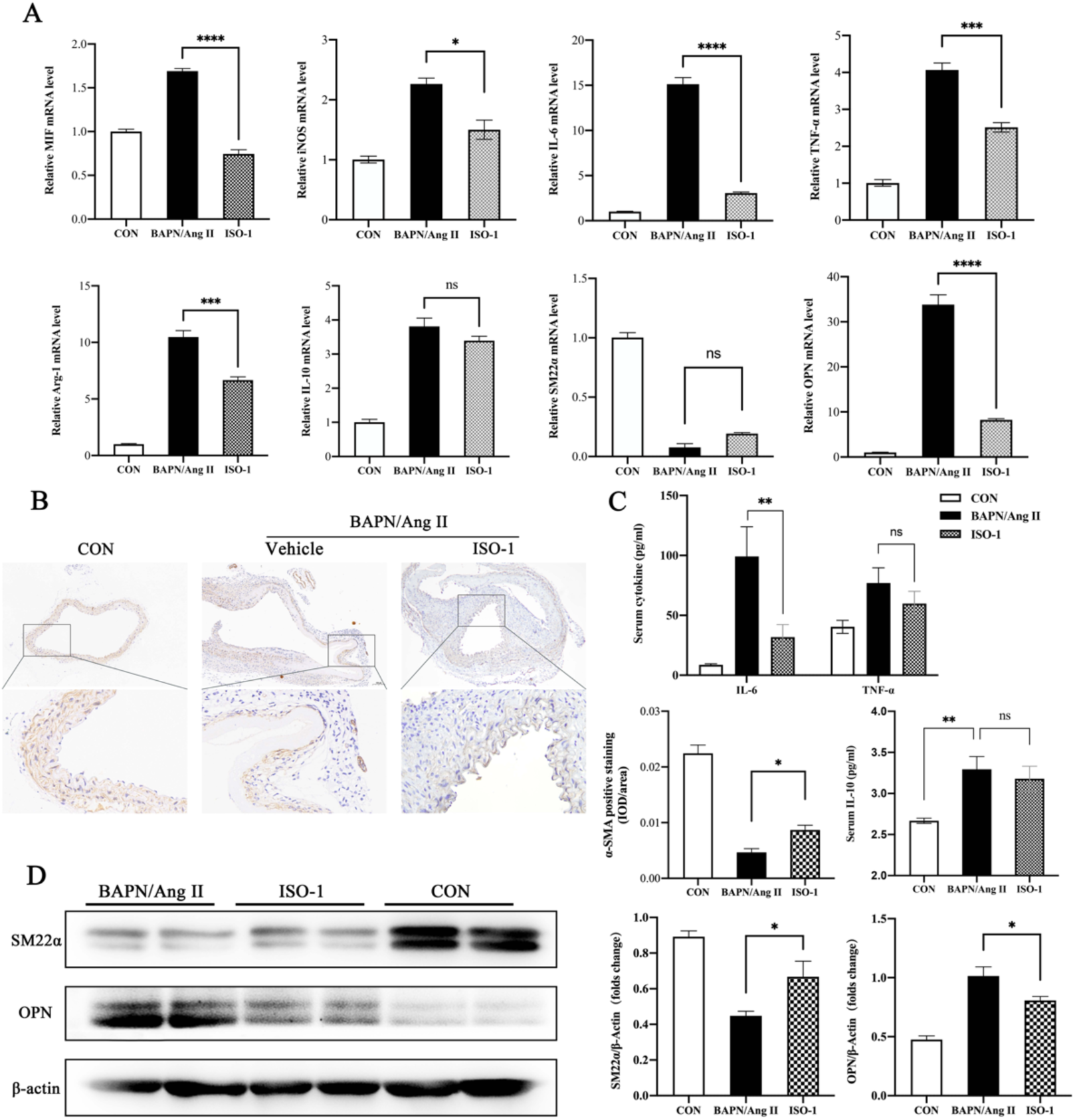
ISO-1 inhibits BAPN/Ang II-induced VSMC phenotypic switching in aorta. **A**, RT-qPCR of MIF, iNOS, IL-6, TNF-α, Arg-1, IL-10, SM22α and OPN in aortic tissues of each group. n=5-7; *P<0.05, ***P<0.001, ****P<0.0001. **B**, ELISA for plasma IL-6, TNF-α and IL-10 levels in mice in each group. **C**, Representative images and quantification of immunohistochemical staining for α-SMA. **D**, Western blot images and quantification of SM22α and OPN in aortic tissues of each group.

### 3. ISO-1 inhibits BAPN/Ang II induced VSMC phenotypic switch

VSMC phenotypic switch is one of the typical characters of AD incidence. We also observed in IHC assays that the expression of the contractile VSMC marker α-SMA was obviously reduced in the middle aorta of mice with aortic dissection, especially at the site of elastic fiber fracture (Fig.3C). The expression of α-SMA in the ISO-1 group was similarly lower than that in the control group but higher than that in the BAPN/Ang II group (P<0.05). Western blot results further proved that the expression of the contractile VSMC marker SM22α protein was significantly decreased and the expression of the synthetic VSMC marker OPN was remarkably increased in the AD group, while ISO-1 antagonized the abnormal expression of SM22α and OPN induced by BAPN/Ang II (Fig.3D)..

Through the above animal experiments, we demonstrated that ISO-1 therapy could further inhibit VSMC phenotypic switch and mid-aortic degradation by attenuating aortic M1 macrophage polarization, thereby alleviating BAPN/Ang II-induced aortic remodeling in mice and reducing aortic dissection morbidity and rupture mortality.

### 4. ISO-1 inhibits LPS/IFN-γ-induced M1 macrophage polarization

In next studies, LPS/IFN-γ stimulation for 24h was established to induce M1 macrophage polarization classically, and RAW264.7 cells were pretreated with different concentrations of ISO-1 prior to stimulation. After pretreatment with ISO-1, protein expression of biomarkers of M1 (iNOS/IL-18) in macrophages were all reduced to various degrees (Fig.4A). The flow cytometry results indicated that the proportion of M1 macrophages (CD86^+^CD206^-^) was significantly elevated to 52.5±3.4% after LPS/IFN-γ stimulation (Fig.4B). In contrast, the proportion of M1 macrophages dropped to 41.4±3.9% after administration of 20 μM ISO-1, demonstrating that inhibition of MIF activity inhibited LPS/IFN-γ-induced M1 macrophage polarization.

**Figure 4.**
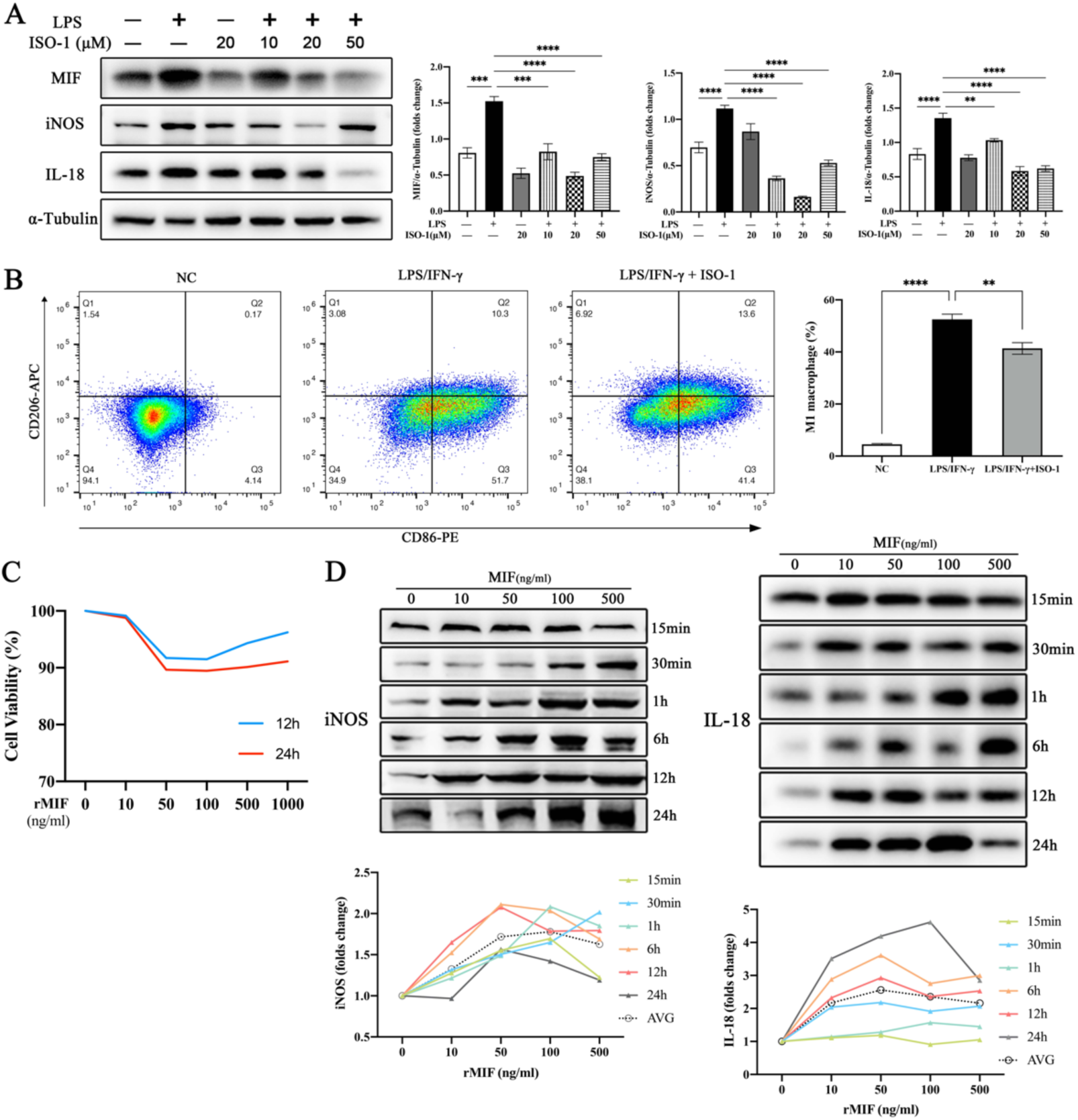
MIF induces polarization of M1 macrophages. RAW264.7 cells pretreated with ISO-1 (10-50μmol/L) and stimulated with LPS/IFN-γ for 24h. **A**, Western blot images and quantification of MIF, iNOS and IL-18 in RAW264.7 cells. **B**, Representative flow cytometry charts of RAW264.7 cells in each group and quantification of M1 macrophages (CD86+CD206-). **C**, Cytotoxic effects of rMIF (0-1000ng/ml) on RAW264.7 cells assessed by CCK8 assay. **D**, Administration of 0-500 ng/ml rMIF stimulated RAW264.7 cells for 15min-24h. Western blot was used to analyze the expression of iNOS and IL-18 proteins, and the protein expression of each group was quantified using the 0ng/ml group as a control.

### 5. Role and mechanism of MIF-mediated polarization of M1 macrophages

We further evaluated the direct effect of exogenous MIF on macrophages. Firstly, we demonstrated by CCK8 assay that recombinant mouse MIF at concentrations ranging from 10-1000 ng/ml had no significant cytotoxicity (Fig.4C). Afterwards, we exposed RAW264.7 cells with a final concentration of 10-500 ng/ml of rMIF for 15 min to 24h. Western blot results revealed that the protein expression levels of iNOS and IL-18 were significantly increased under rMIF stimulation at concentrations of 50 to 100 ng/ml and were time-dependent (Fig.4D), with the strongest expression at 24 h of stimulation (Fig.5A). RT-qPCR results also indicated that the mRNA expression levels of IL-6, TNF-α and iNOS were significantly higher under the above conditions of stimulation compared with the control group (Fig.5B). The cell morphology and intracellular iNOS expression of each group were detected by IF, and it was found that macrophages induced by LPS/IFN-γ had omelet-like morphology and significantly higher expression of iNOS in the cell cytoplasm, while some cells in the MIF group were also omelet-like with short tentacles, and the intracytoplasmic iNOS fluorescence intensity was significantly higher than that of the control group (P<0.05) (Fig.5C). Flow cytometry data showed a significant increase in the proportion of M1 macrophages after 50 and 100 ng/ml rMIF stimulation for 24h compared to the control group (Fig.5D). The concentration of pro-inflammatory factors in the culture medium was measured by ELISA, suggesting that the secretion of IL-1β, IL-6 and TNF-α was elevated after rMIF induction at 50-100 ng/ml, with IL-6 and TNF-α being elevated to a greater extent (Fig.6A). Taken together, these results proved that MIF induced M1 polarization in RAW264.7 cells, as evidenced by elevated expression of M1 macrophage-specific markers, antibodies, and enhanced secretion of pro-inflammatory factors.

**Figure 5.**
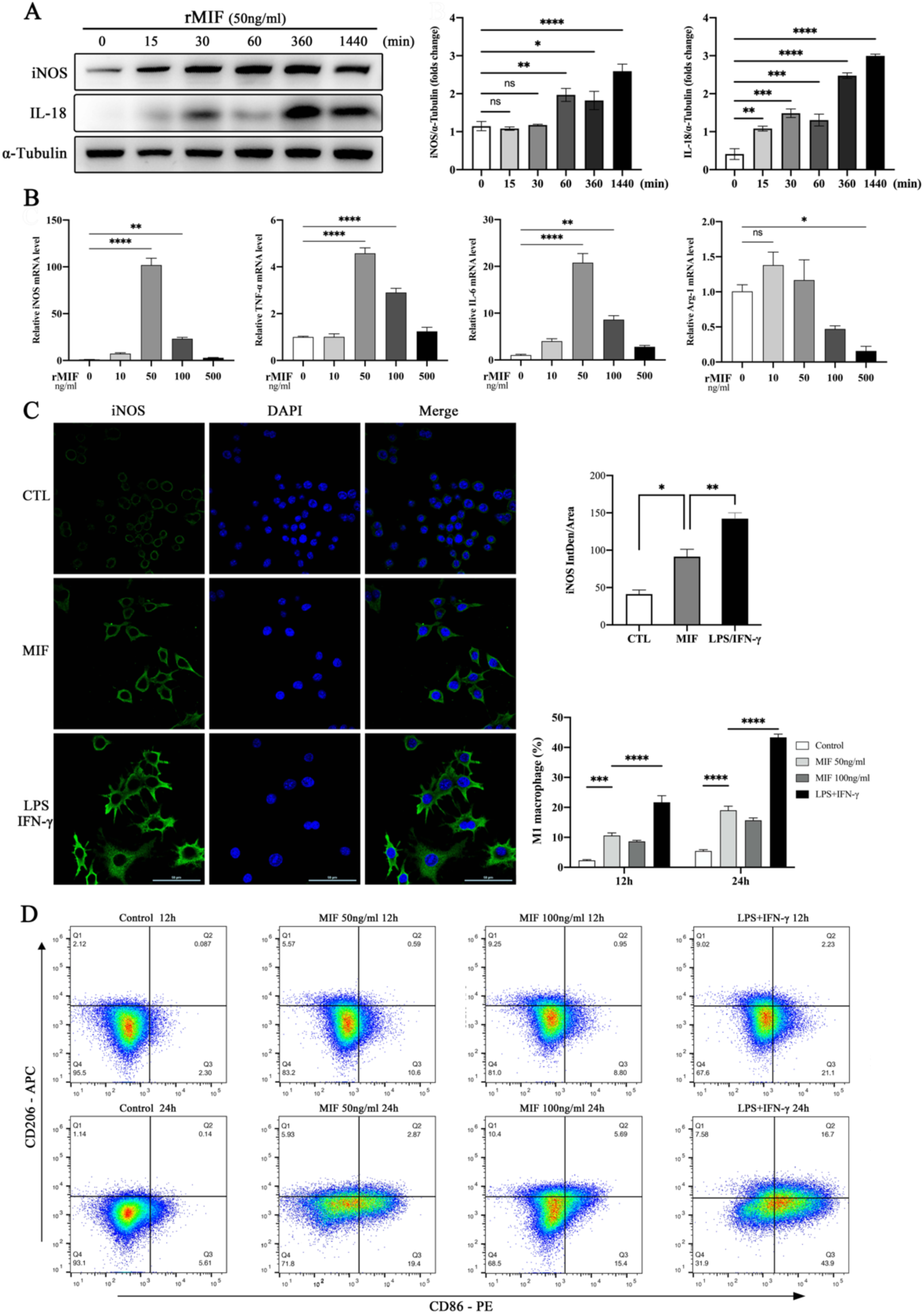
MIF induces polarization of M1 macrophages. **A**, Western blot images and quantification of iNOS and IL-18 in RAW264.7 cells treated by 50ng rMIF for 15min-24h. **B**, RT-qPCR of iNOS, TNF-α, IL-6 and Arg-1 in RAW264.7 cells treated by 0-500 ng/ml rMIF. **C**, Representative photographs and quantification of immunofluorescence staining of iNOS (green) and DAPI (blue) in RAW264.7 cells treated with MIF or LPS/IFN-γ. **D**, Representative flow cytometry charts of RAW264.7 cells treated with MIF or LPS/IFN-γ and quantification of M1 macrophages (CD86+CD206-).

**Figure 6.**
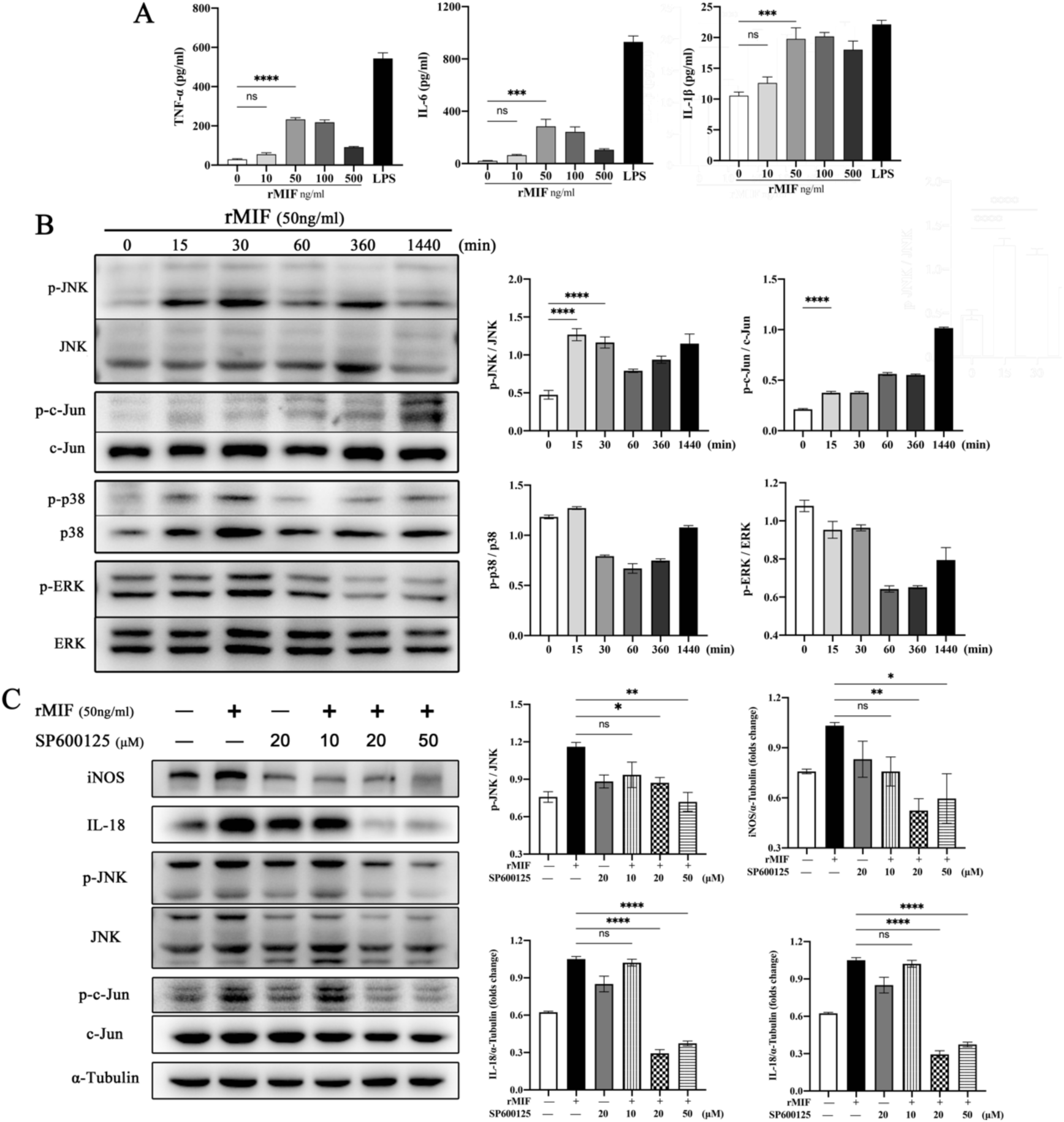
MIF stimulates M1 macrophage polarization via the JNK/c-Jun pathway. **A**, Measurement of IL-6, TNF-α and IL-1β concentrations in the cell culture medium of each group by ELISA. **B**, Western blot images and quantification of JNK, c-Jun, p38, and ERK phosphorylation after different lengths of MIF stimulation. **C**, Western blot images and quantification of iNOS, IL-18 protein expression and JNK, c-Jun phosphorylation in RAW264.7 cells pretreated with SP600125 after MIF stimulation.

### 6. MIF-induced M1 macrophage polarization depends on the JNK/c-Jun pathway

To investigate whether MIF-induced macrophage polarization was mediated by the MAPK pathway, we tested the phosphorylation of ERK1/2, P38 and JNK after rMIF treatment (Fig.6B). Upregulation of JNK phosphorylation was observed 15 min after exposure of RAW264.7 cells with 50 ng/ml of rMIF, which was approximately 1.85-fold higher than that of the control group and lasted for 24 h. In contrast, phosphorylation of P38 and ERK was not found to be sequentially enhanced.

Correspondingly, we also assessed the level of phosphorylation of c-Jun, a downstream substrate of JNK, and could observe that c-Jun activation roughly paralleled JNK, with enhanced phosphorylation observed at 15 min, reaching a peak phosphorylation at 30 min and lasting approximately 24 h. To verify whether MIF-mediated M1 polarization is dependent on the JNK/c-Jun pathway, we pretreated RAW264.7 cells with the JNK inhibitor SP600125 for 30 min prior to rMIF stimulation. Western blot showed that SP600126 successfully blocked JNK/c-Jun pathway phosphorylation and markedly reduced the protein expression of M1 macrophage markers (P<0.01) (Fig.6C).

### 7. MIF-targeted CD74/CXCR2 membrane receptor complex induces M1 polarization

We found that the mRNA expression of CD74, CXCR2 and CXR4 in RAW264.7 cells stimulated by 50ng/ml of rMIF was increased to different degrees by RT-qPCR assay (Fig.7A). To further clarify the pathway of MIF-activated JNK phosphorylation, we first used siRNA to interfere with CD74 gene expression. After confirming the downregulation of CD74 protein expression in macrophages, cells were stimulated using rMIF under the above conditions (Fig.7B). The phosphorylation of JNK and c-Jun was inhibited by antagonizing CD74, and the protein expression of iNOS and IL-18 was significantly lower than that of the unantagonized group (Fig.7C).

**Figure 7.**
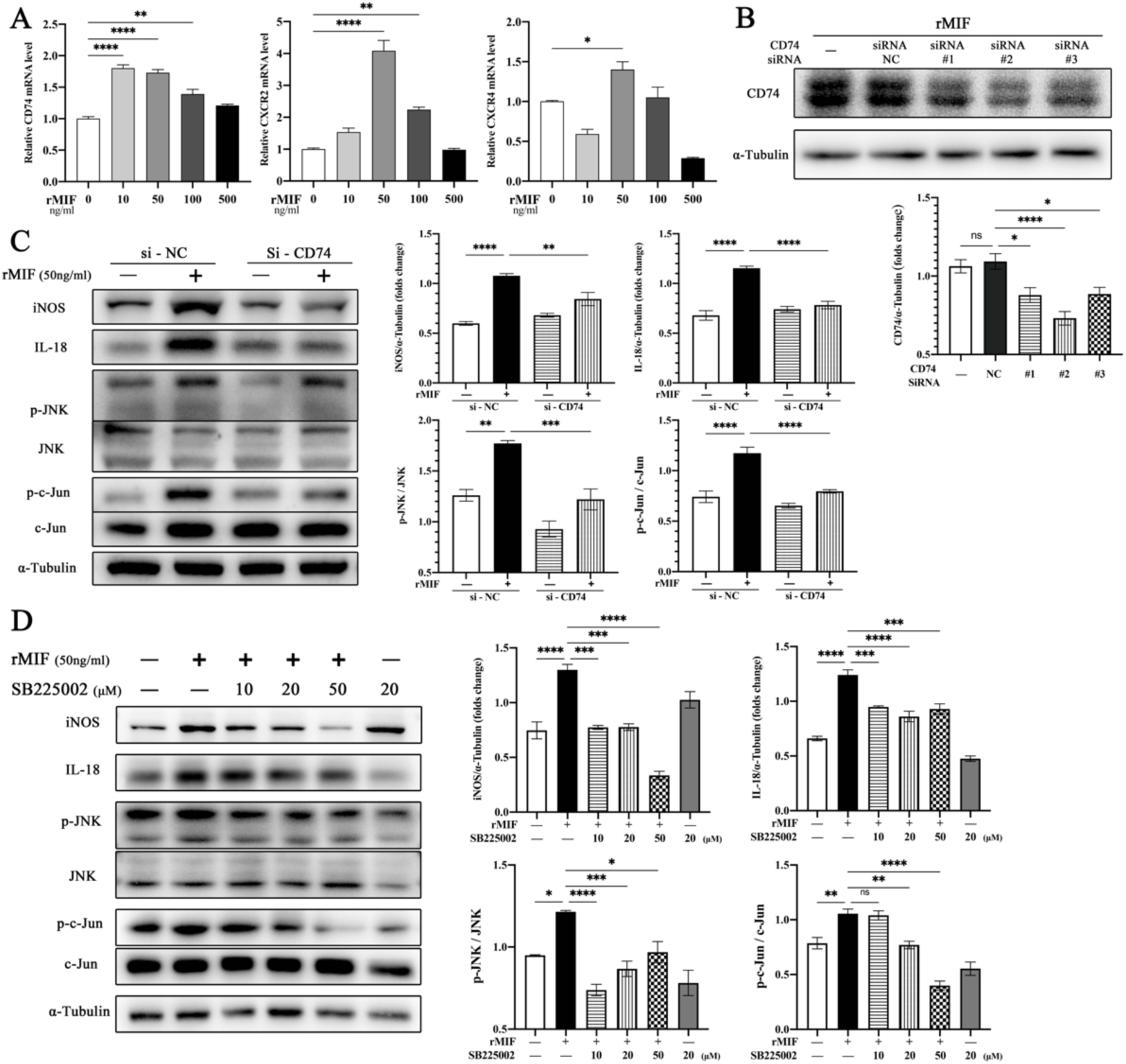
MIF-induced M1 macrophage polarization depends on CD74/CXCR2. **A**, RT-qPCR of CD74, CXCR2 and CXCR4 in RAW264.7 cells treated by 0-500 ng/ml rMIF. **B**, Western blot assessment of CD74 siRNA silencing levels in RAW264.7 cells. **C**-**D**, Western blot images and quantification of iNOS, IL-18 protein expression and JNK, c-Jun phosphorylation in RAW264.7 cells pretreated with siRNA-CD74 (C) or SB225002 (D) after MIF stimulation.

Similarly, we pretreated RAW264.7 cells with the pharmacological CXCR2 antagonist SB225002 and the CXCR4 antagonist AMD3100, respectively. Both JNK/c-Jun pathway activation and M1 marker protein expression were blocked in RAW264.7 cells after antagonism of CXCR2, while similar effects were not seen with CXCR4 inhibition (Fig.7D). Integrating this part of the cell experiments, we demonstrated that MIF could promote M1 polarization in RAW264.7 cells by targeting the CD74/CXCR2 receptor complex and activating the intracellular JNK/c-Jun signaling pathway.

### 8. Co-culture with M1 macrophages induces VSMC phenotypic switch and apoptosis

To simulate the pathological process of aortic dissection, we established an indirect co-culture system of RAW264.7 cells with mouse aortic smooth muscle (MOVAS) cells. IF results showed a significant reduction in the fluorescence intensity of the contractile marker SM22α in MOVAS cells after co-culture with macrophages that had undergone MIF or LPS exposure, whereas that of the synthetic marker OPN was significantly elevated (Fig.8A). We got similar outcomes by WB assay, the MOVAS cells in LPS and MIF groups underwent phenotypic switching, the protein expression of contractile markers SM22α and MYH11 were significantly reduced compared with the control group, while the protein expression of OPN was significantly increased, and the extent of phenotypic switch was stronger in the LPS group than in the MIF group (Fig.8B). Subsequently, we clarified the apoptosis of MOVAS cells in each group by flow cytometry, and it was observed that the proportion of apoptotic cells in MOVAS cells after co-culture in the MIF and LPS groups was significantly increased, and it was more in the LPS group than in the MIF group (Fig.9A).

**Figure 8.**
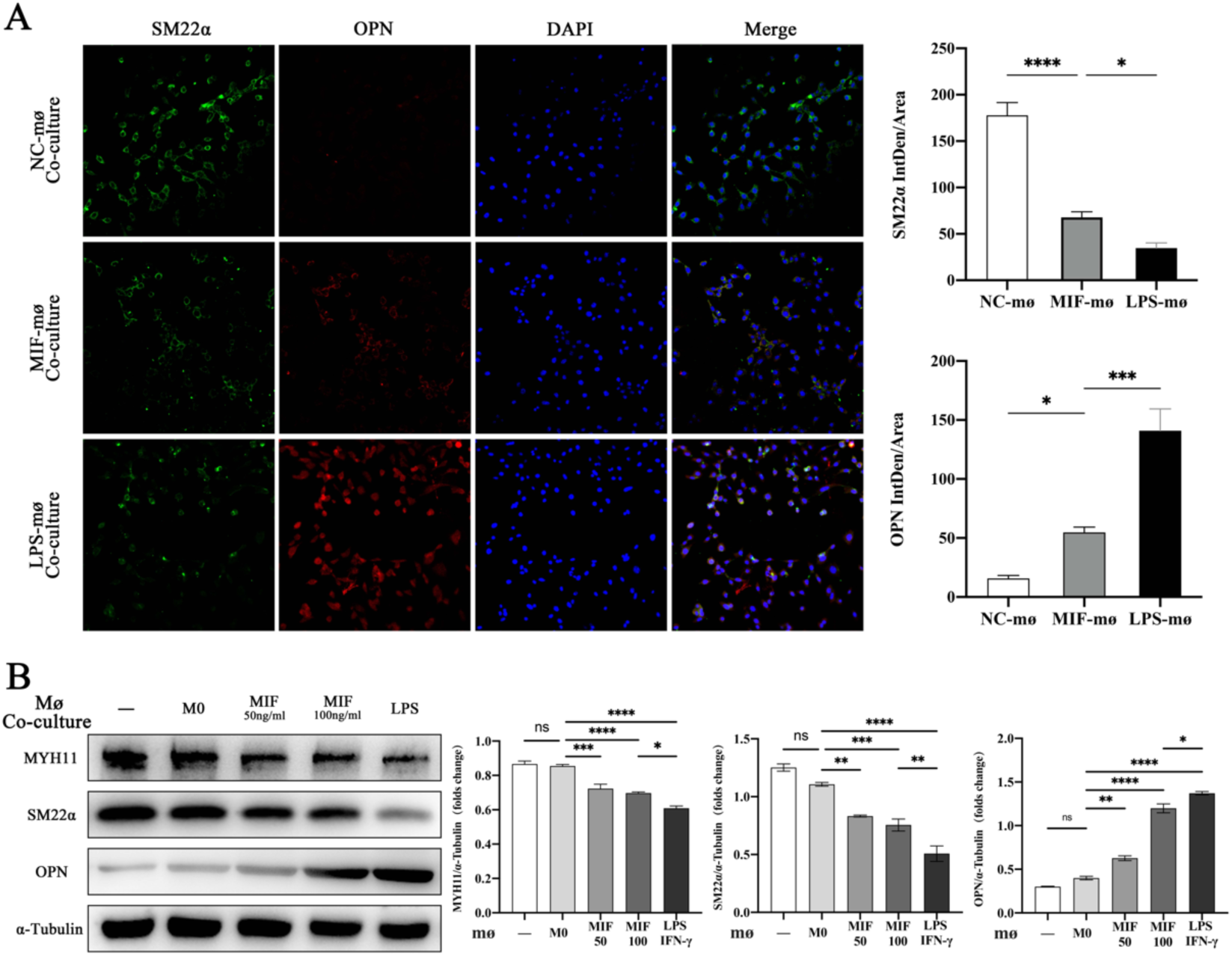
Co-culture with M1 macrophages accentuates MOVAS cell phenotype switching. RAW264.7 cells were stimulated with various polarizations and then indirectly co-cultured with MOVAS cells for 48h. **A**, Representative photographs and quantification of immunofluorescence staining of SM22α (green), OPN (red) and DAPI (blue) in MOVAS cells of each group. **B**, Western blot images and quantification of MYH11, SM22α and OPN in MOVAS cells co-cultured with RAW264.7 cells for 48h. **C**, Gelatin gel zymography of MMPs activity in MOVAS cells.

**Figure 9.**
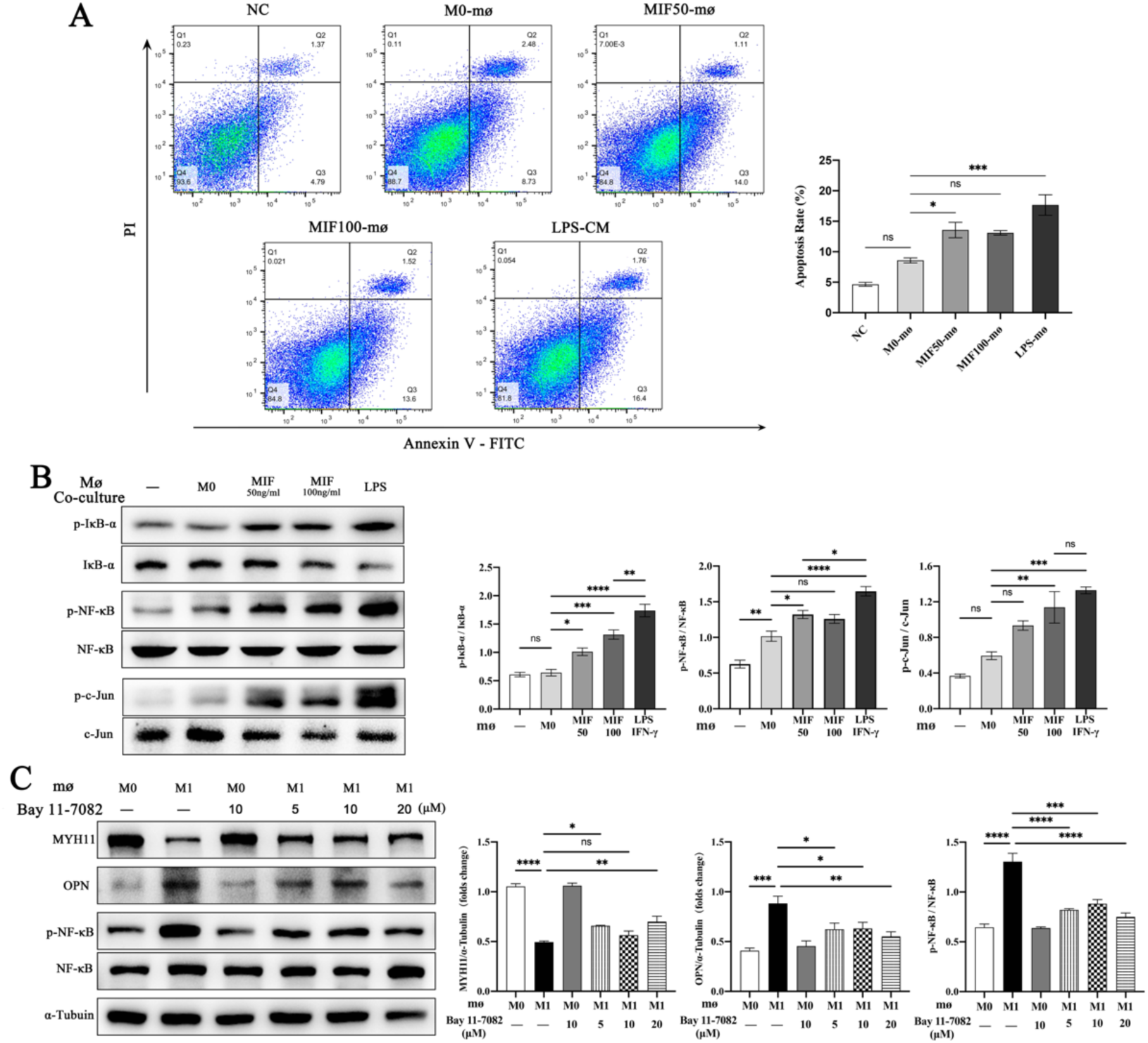
M1 macrophages mediate MOVAS cell phenotype switching via the NF-κB signaling pathway. **A**, Representative flow cytometry charts of MOVAS cells co-cultured with RAW264.7 cells for 48h and quantification of apoptotic MOVAS cells (Annexin V+PI-). **B**, Western blot images and quantification of NF-κB, IκB-α and c-Jun phosphorylation in MOVAS cells of each group. **C**, MOVAS cells were pretreated with BAY 11-7082 and co-cultured with macrophages for 48 h. MYH11 and OPN protein expression and NF-κB phosphorylation levels were assessed by western blot.

We noticed enhanced phosphorylation of NF-κB and c-Jun in MOVAS cells after co-culture with M1 macrophages (Fig.9B), so we further pretreated MOVAS cells with the NF-κB inhibitor BAY 11-7082 and the c-Jun inhibitor T-5224, respectively, prior to co-culture. After pretreatment with BAY 11-7082, the activation of NF-κB in MOVAS cells was inhibited and the aberrant expression of phenotypic switching markers caused by co-culture with M1 macrophages was reversed (Fig.9C). In contrast, no apparent changes in phenotypic switching markers were observed after pretreatment with T-5224. As demonstrated in this part of the experiment, after co-culture with MIF or LPS/IFN-γ-induced M1 macrophages, the NF-κB pathway was activated in MOVAS cells, which induced phenotypic switch and apoptosis in MOVAS cells.

## Discussion

In this research, we conducted a basic study by establishing a mouse model of acute aortic dissection. After integrating previous studies^24–26^ and pre-testing the modeling conditions, we determined a way to establish the acute aortic dissection model in mouse using a concentration of 0.5% BAPN drinking water continuously fed to mice for 2 weeks, followed by two subcutaneous injections of Ang II at a concentration of 0.72 mg/kg. The incidence of AD in this group reached 86.7% in the pre-trial, with a rupture rate of 40%. Macrophage migration inhibitory factor (MIF) is a pro-inflammatory chemokine-like cytokine that has been reported to mediate a variety of inflammatory and neoplastic diseases through macrophage polarization and smooth muscle cell differentiation.^21,27,28^ To investigate whether MIF is involved in the pathogenesis of aortic dissection by modulating macrophage polarization, we administered ISO-1, a MIF-specific inhibitor, to antagonize MIF activity on the basis of the above mouse aortic dissection model.

During the experiment, we noticed that administration of ISO-1 was effective in slowing down BAPN/Ang II-induced aortic remodeling and reducing the occurrence of aortic dissection. We also demonstrated that in aortic dissection, massive M1 macrophages infiltrate the outer and middle layers of the aorta, and the ISO-1 can suppress the expression of M1 macrophage markers and the secretion of pro-inflammatory factors in the aorta. VSMC phenotypic switch refers to the pathological stimulation of contractile VSMCs to de-differentiate into synthetic VSMCS, which accelerates aortic remodeling and leads to progressive aortic dilatation and rupture.^29^ In the present study, we confirmed that VSMC phenotypic switch occurred in the aorta of BAPN/Ang II-induced mice and that the VSMC phenotypic switching process was inhibited in the ISO-1 group. Activation of the inflammatory system is the initiating factor of aortic dissection and is also present throughout the development of AD. By performing studies in mice, we demonstrated that pharmacological inhibition of MIF attenuates VSMC phenotypic switch and the extent of mid-aortic degradation by inhibiting M1 macrophage polarization, infiltration, and associated inflammatory factor secretion, thereby slowing BAPN/Ang II-induced aortic remodeling and the development of aortic dissection in mice.

In order to make clear whether MIF gets involved in the regulation of macrophage polarization, we confirmed by a series of in vitro experiments that ISO-1 can effectively inhibit the activity of MIF in LPS/IFN-γ-induced M1 macrophages and inhibit the expression of specific markers and surface antibodies of M1 macrophages. Next, we also found that 50-100ng/ml rMIF can target CD74/CXCR2 receptor complex on RAW264.7 cell membrane, and up-regulate the expressions of specific markers and surface antibodies and the secretion of pro-inflammatory factors of M1 macrophages by activating JNK/c-Jun/AP-1 signaling pathway in cells, thereby polarizing macrophages to M1. CD74 is a non-polymorphic transmembrane protein and a high-affinity receptor of the cell membrane of MIF family, which facilitates the synthesis and secretion of pro-inflammatory factors and cellular adhesion molecules and mediates the occurrence of a variety of inflammations and immune diseases.^30,31^ Studies have shown that administrating CD74 shRNA intravenously can inhibit M1 polarization and insulin resistance of macrophages in adipose tissues of mice induced by high fat diet.^32^ An opposite trend, however, was observed in tumors. The expression of CD74 mRNA in patients with brain tumor increased with malignancy degree worse. Brain tumor cells can target CD74 by secreting MIF, promote microglial cells to polarize from M1 to carcinogenic M2, inhibit the secretion of IFN-γ in microglial cells, and get involved in the immune escape mechanism of tumor.^19,33^ In addition to CD74, MIF has also been identified to be a non-homologous ligand of CXCR2 and CXCR4. Extracellular MIF binds to the cell membrane receptor CD74 and forms signaling complexes with the G-protein-coupled chemokine receptors CXCR2 and CXCR4, which mediate the MIF-associated signaling pathway in addition to the signaling pathways of their cognate ligands CXCL8 and CXCL12, respectively.^34^ The expression of CXCR2 is mainly limited to monocytes and neutrophils,^35^ while CXCR4 is widely expressed in hematopoietic cell lines, endothelial cell lines and nerve cells.^36,37^ MIF/CXCR2/CXCR4 axis can promote atherosclerosis, widely stimulate vasculitic response and promote the recruitment of monocytes and neutrophils through CXCR2, and target the recruitment of T cells via CXCR4, promote acute and chronic inflammation through the recruitment of CXCR4 targeting T cells, the rapid activation of integrin and the influx of calcium ions.^38,39^

In the investigation of intracellular signaling pathway, we detected the phosphorylation levels of three pathways in MAPK family in macrophages stimulated by rMIF, and only observed the transient and sustained phosphorylation of JNK. The downstream c-Jun was also significantly phosphorylated, which was synchronous with the phosphorylation of JNK. After phosphorylation, c-Jun translocated into the nucleus, and the activation of the nuclear AP-1 led to the increase of mRNA expression of pro-inflammatory gene, as well as the production of pro-inflammatory cytokines and chemerins, including TNF-α, IL-6, COX-2 and CCL-12, etc.^40^ The expression and secretion of MMP9 also depended on the activity of AP-1.^41^ In macrophages, LPS may initiate MAPK signaling cascade via TLR4/MyD88, activate AP-1 in the nucleus, and contribute to the release of pro-inflammatory factor.^42^ As the core component of AP-1, c-Jun family was particularly crucial in the polarization process of macrophages. C-Jun can perceive a wide range of macrophage activation signals, including Th1 and Th2 cytokines and pro-inflammatory and anti-inflammatory stimuli. C-Jun maintained the occurrence of inflammation by increasing the expression of pro-inflammatory genes, for example COX-2, and inhibiting the expression of M2 macrophage marker Arg-1, which got involved in inflammation resolution and tissue repair.^43^ Thus, JNK/c-Jun/AP-1 pathway has been widely studied as an anti-inflammatory target. In fibroblasts and T cells, it was observed that MIF triggered the transient activation of JNK/c-Jun/AP-1 pathway via the upstream CD74/CXCR4/SRC/PI3K axis and can induce CXCL8 secretion and rapidly expedite the inflammatory process.^44^ In this study, it was confirmed that MIF could target CD74/CXCR2 receptor complex, and macrophages underwent M1 polarization, as induced by JNK/c-Jun/AP-1 signaling pathway.

The immediate cause of the formation of aortic dissection is the destruction of the integrity of aortic structure and the decrease of arterial compliance. Since the supporting power of aorta is mainly provided by the middle smooth muscle layer, ECM degradation and VSMC apoptosis can weaken the aortic wall, the aorta lost the integrity of overall structure and intrinsic molecular connectivity, which ultimately lead to aortic dissection. As the key node of AD, the phenotypic switch of VSMC is also the pathological basis of atherosclerosis, vascular calcification, vascular aging and hypertension.^45^ Aortic dissection is a complex pathological process jointly participated in by a variety of immune cells, smooth muscle cells and endothelial cells. Intercellular interaction and cell-ECM interaction played a non-negligible part in the pathogenesis of AD.

We have noticed some limitations of this study. The interaction between macrophages, as the most significantly infiltrated immune cells in the early stage of AD, and smooth muscle cells may directly influence the progress of dissection. Both smooth muscle cells and endothelial cells can express intercellular adhesion molecule (ICAM-1), vascular cell adhesion molecule-1 (VCAM-1) and monocyte chemoattractant protein-1 (MCP-1) recruit macrophages to infiltrate aorta, and aggravate chronic inflammation.^46,47^ The activated inflammatory macrophages may up-regulate the expression of Fas protein in VSMC via TNF-α and IL-1β, promote VSMC apoptosis, and the proliferation ability of VSMC was also influenced by macrophages, which regulated the formation of intima.^48,49^ Through single cell sequencing, it was found that the receptor-ligand interaction between VSMC and immune cell population in aortic dissection was significantly up-regulated, especially for macrophages and T cells. Therefore, compared with single cells such as macrophages or VSMCs, the interaction between two or even more types of cells played a more significant role in the process of vascular remodeling in aortic dissection.^50^ In our study, we observed that inhibiting the infiltration and polarization of M1 macrophages in aortic tissues can weaken the phenotypic switch of VSMCs and lower the vascular remodeling of aorta. Therefore, the interaction between M1 macrophages and VSMCs may be built on paracrine and play a promoting role in phenotypic switch of VSMCs. To verify this speculation, by establishing a co-culture system for M1 macrophages and mouse aortic smooth muscle cells with different polarization degrees, we observed that MOVAS cells co-cultured with M0 macrophages didn’t undergo significant phenotypic switch. After being co-cultured with M1 macrophages, MOVAS cells underwent significant phenotypic switch and apoptosis. At the same time, we also observed the phosphorylation expressions of NF-κB and c-Jun, key nodes in TNF-α and IL-6 signaling pathways. Subsequent experiments documented that abnormal phenotypic switch can be redressed by treating MOVAS with the NF-κB antagonist, but similar results were not got with the c-Jun antagonist which proved that NF-κB was a key factor of phenotypic switch of VSMCs induced by M1 macrophages.

Recent studies have shown that with high plasticity, VSMC has multiple differentiation potentials in different disease states and eventually transform into abnormal phenotypes of morphological and functional changes.^51,52^ There are multiform phenotypes in aortic aneurysm specimens, such as contractile VSMC, mesenchymal-like VSMC, macrophage-like VSMC and fibroblast-like VSMC.^53^ Among them, mesenchymal-like VSMC and fibroblast-like VSMC probably play a role in repairing aortic dissection and aortic aneurysm, while macrophage-like VSMC not only has phagocytic function, but also interacts positively with immune cells, recruits monocytes and lymphocytes, stimulate them to migrate to aortic wall and exert their pro-inflammatory function.^54^ In this study, M1 macrophages may induce phenotypic transformation of MOVAS cells, but it is not verified whether this is associated with a variety of phenotypes such as macrophage-like VSMC and fibroblast-like VSMC.

There are still many limitations in this study. Firstly, in the animal experiment, we were unable to evaluate the effect of MIF overexpression on BAPN/Ang II-induced aortic dissection in mice. Secondly, we proved the role and pathway of MIF-mediated macrophage polarization but failed to identify whether this polarization was based on the autocrine effect of macrophages or the paracrine effect of other cells. Finally, this study didn’t specify which cytokine in M1 macrophages mediated the phenotypic switch of VSMCs, which warranted further study.

To sum up, we observe the regulatory role that MIF plays in macrophage polarization and VSMC phenotypic switch in the aortic dissection model at the animal level. The promotion of MIF on M1 macrophage polarization and the M1 macrophage-mediated VSMC phenotypic switching process were also confirmed at the cellular level. We tie two key pathological characteristics of AD together, macrophage polarization and VSMC phenotypic switch, rather than simply focus on a given node. This study provides a new train of thought for aortic dissection and other vascular diseases with similar pathological processes. The elaboration of the regulatory role and mechanism of MIF in the progression of aortic dissection may deliver a new target and strategy for early diagnosis, prevention, and treatment of aortic dissection.

